# The strength of weird ties: positive genetic interactions have greater scientific impact but are under-represented in the literature

**DOI:** 10.1101/2025.06.25.661621

**Authors:** Mengyi Sun, David M. McCandlish

## Abstract

Genetic interactions, where the combined effect of perturbing two genes leads to a phenotype that deviates from the expectation based on the effects of the individual perturbations, often provide important information about the functional architecture of biological systems. These interactions are commonly classified as negative or positive based on whether the phenotype of the double mutant is less than or greater than what would be expected based on the single mutant effects. When the trait in question is fitness, systematic studies of pairwise deletions have shown that negative interactions typically link genes with similar functional annotations, while positive interactions typically link genes that are less obviously related and thus often viewed as less informative. However, research in the sociology of science suggests that transformative discoveries often arise from unexpected, cross-domain linkages. To evaluate these competing perspectives, we integrated large-scale genetic interaction data in yeast with literature annotations from the Saccharomyces Genome Database and citation data from iCite. We found that positive genetic interactions are associated with greater scientific impact, as measured by citation metrics, contrary to prevailing assumptions. Despite their greater impact, positive interactions are significantly underrepresented in the scientific literature. Interestingly, the advent of an unbiased mapping technology capable of detecting positive and negative interactions with equal facility did not alleviate this bias, implicating factors beyond mere discoverability. We also find that the research building on positive interactions forms a more interconnected body of follow-on work. Together, our analyses reveal the underappreciated value of positive genetic interactions and suggest that investigating cross-domain genetic links represents an under-exploited opportunity deserving of greater attention.

## Introduction

Genetic interaction refers to the phenomenon where the combined effect of disrupting two genes on a trait deviates from the expectation based on perturbing each gene individually^1^. This phenomenon has proved valuable for uncovering functional relationships among genes and pathways^2^, and has implications for a range of biological questions: from the origin of species in evolutionary biology^3^ to the genetic basis of human diseases^4^ and drug interactions^5^.

Concretely, in a model organism such as yeast, researchers can disable (‘knock out’) individual genes and measure the effect on fitness, defined here as how fast a population of cells grows. A genetic interaction asks whether disabling two genes together has the effect one would predict from disabling each gene on its own. If the double knockout grows worse than predicted, the two genes have a negative interaction; the canonical example is known as synthetic lethality^6^, in which a cell tolerates the loss of either gene by itself but dies when both are lost. If the double knockout instead grows better than predicted, the interaction is positive. The interaction score quantifies this deviation from the independent-effects prediction: writing f_a_ and f_b_ for the fitnesses of the two single-gene knockouts and f_ab_ for that of the double knockout, the score is I_ab_ = f_ab_ − (f_a_ × f_b_), and its sign (negative or positive) is the property we study.

Starting in the early 2000s, Synthetic Genetic Array (SGA) technology made it possible to measure interactions of both signs across the entire yeast genome^2,7,8^. In an SGA experiment, a strain carrying one gene deletion is systematically mated to an ordered array of strains that each carry a different single-gene deletion; this produces, for thousands of gene pairs, cells in which both genes are deleted, whose growth is then measured and compared with the single-deletion effects to score the interaction. Prior studies^9^ have shown that negative interactions tend to arise between genes that are already known to be functionally related, that is, genes participating in the same or closely connected biological processes, so that negative interactions are widely regarded as informative about how genes work together^9^. Positive interactions, which tend to link genes without apparent functional relationship, are often deemed less informative^9^. This perspective implies that studying gene pairs involved in negative interactions should be scientifically more valuable.

However, a key insight from the sociology of science is that impactful innovation often stems from unexpected linkages across conceptual domains^10^. From this perspective, positive genetic interactions, by linking genes whose functional roles are less obviously related, may offer greater potential for scientific innovation.

To evaluate whether positive or negative interactions are more scientifically impactful, we analyzed large-scale genetic interaction data from yeast, linking gene pairs to their associated scientific output using yeast gene literature^11^ and iCite citation databases^12^. Our results show that positive interactions are associated with greater scientific impact. Nevertheless, despite their empirical importance, positive interactions remain significantly underrepresented in the scientific literature. Interestingly, the introduction of SGA, an unbiased mapping technology that detects positive and negative interactions with equal facility, did not relieve this under-representation but instead deepened it relative to the prior trend, indicating that factors beyond experimental discoverability are at play. These findings challenge conventional views about functional informativeness and highlight the value of applying sociology-of-science insights to biological discovery.

## Results

### Positive genetic interactions are associated with higher scientific impact

We investigated the scientific impact of positive versus negative gene interactions by analyzing publications from the Saccharomyces Genome Database (SGD)^11^. This database links 418,111 papers to their associated genes, from which we selected the subset of publications associated with exactly two genes. We then further selected the subset of these papers where the gene pair studied exhibited a significant genetic interaction score (p < 0.05) from the large-scale screen conducted by Costanzo et al.^2^. To assess scientific impact, we used iCite, a bibliometric database accessed via PubMed IDs provided by SGD. Our primary metric was the year-specific percentile citation rank, which compares a paper’s citation count to those of other papers published in the same year within the SGD corpus. This approach standardizes citation impact by controlling for publication year (a standard practice in scientometrics) to avoid biasing results in favor of older papers that have had more time to accumulate citations^13^.

As shown in Figure 1A, publications focusing on positive genetic interactions are significantly more impactful than those involving negative interactions (p < 0.01, t-test; p < 0.01, Mann-Whitney U test), as measured by year-specific percentile citation rank. This finding is further supported by the year-normalized citation impact^14^ (a paper’s citation count divided by the average for papers published in the same year), where Figure 1B shows that studies on positive interactions tend to have ∼20% more citations on average than those on negative interactions of the same age (p < 0.01, t-test; p < 0.01, Mann-Whitney U test). These results consistently indicate that when gene pairs involved in positive interactions are studied together, they tend to generate more impactful research, as quantified by this observed citation premium.

**Figure 1.**
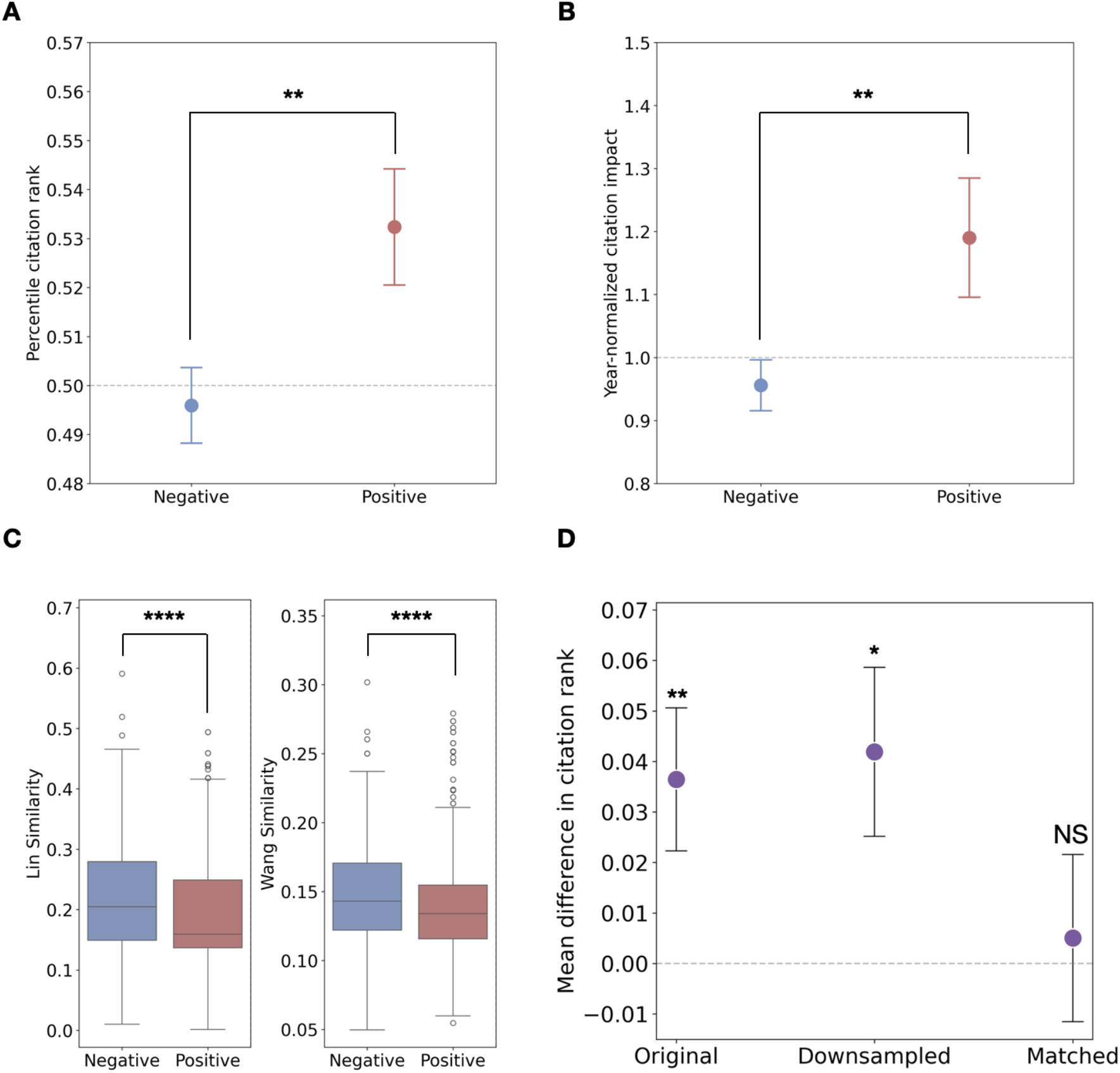
Positive interactions are linked to higher scientific impact and novelty. (A) Mean year-specific percentile citation rank and (B) year-normalized citation impact for literature on positive (red) versus negative (blue) interactions. (C) GO similarity between the paired genes, measured by Lin’s method (left) and Wang’s method (right), for positive (red) versus negative (blue) interactions. (D) Mean difference in citation rank (positive − negative) across the original, downsampled, and GO-similarity-matched samples; the citation advantage of positive interactions vanishes only after matching on GO similarity. All error bars represent standard errors. Significance levels: NS, P ≥ 0.05; *P < 0.05; **P < 0.01; ***P < 0.001; ****P < 0.0001.

We further explored whether positive interactions’ higher impact stems from their novelty, i.e. their propensity to link apparently disparate biological processes. Figure 1C confirms positive interaction pairs consistently exhibit lower GO similarity than negative interactions (using Lin’s^15^ and Wang’s^16^ metrics, all p < 0.0001, Mann-Whitney U test), indicating greater novelty. To confirm novelty as the mediator, we analyzed citation rank differences across sample types (Figure 1D). The full dataset shows a significant citation advantage for positive interactions (p < 0.01, t-test, Figure 1D, left). Crucially, when we match positive and negative interaction pairs based on their GO similarities (annotations propagated to all parent terms, each positive pair matched to a negative pair by Mahalanobis distance on its (Wang, Lin) similarity vector, where lower similarity reflects greater novelty; see Methods), this advantage becomes statistically indistinguishable from zero (p = 0.8, t-test, Figure 1D, right). This indicates novelty drives the observed citation advantage. To confirm that the loss of statistical significance was not a sample size artifact, we also performed random downsampling to equalize positive and negative interaction paper counts. In this downsampled scenario, the citation advantage for positive interactions remained significantly higher than zero (p < 0.05, t-test, Figure 1D, middle), similar to the full sample.

These conclusions are robust to the specific analytical choices made above. In a multivariate regression framework (Table S1; ordinary linear regression), the positive-interaction citation premium remains positive after controlling for the number of authors, journal venue, publication year, and the paper’s bibliometric subfield (its OpenAlex primary topic^17^, which—unlike Gene Ontology annotations—is derived from the paper’s text and citations and so provides a gene-independent control for field composition). The premium also stays statistically significant across the entire control ladder for the percentile citation rank; for year-normalized citations it remains significant through the confounder-adjusted model and becomes marginal only in the most saturated specification (p = 0.06), and controlling for the interaction magnitude |I| if anything further strengthens the estimated citation premium in favor of positive interactions. The mediation by novelty is also not specific to Gene Ontology annotations: when we instead measure functional similarity by KEGG pathway co-membership^18^, positive interactions again link genes that share fewer pathways, and matching positive to negative pairs on this non-GO measure renders the citation-rank advantage statistically non-significant (matched p = 0.12, versus p < 0.01 before matching), mirroring the GO-similarity result and indicating mediation by novelty rather than by any particularity of GO annotation. These additional robustness checks indicate that the citation premium is not an artifact of a particular confounder or the specifics of GO annotation practices.

Together, these findings suggest that gene pairs involved in positive genetic interactions are associated with higher-impact publications because positive genetic interactions often uncover novel connections between seemingly unrelated biological processes.

### Positive genetic interactions are underexplored in the scientific literature

Because gene-pairs exhibiting positive interactions tend to be associated with higher impact, we wondered if positively interacting gene pairs are also enriched in literature. We compared the fraction of positive and negative gene pairs in the literature versus those identified through unbiased experimental screening. To our surprise, positively interacting gene pairs are significantly underrepresented in the literature compared to their prevalence in unbiased experimental screens (p < 0.0001, Fisher’s exact test, Figure 2A). For instance, while approximately 48% of measured gene pairs exhibit positive interactions, among co-studied gene pairs that are involved in significant genetic interaction, only 30% are positive interactions. This depletion suggests a systematic bias against studying positive interactions. This underrepresentation is further highlighted by Figure 2B, which plots the log odds ratio of positive to negative interactions in the literature compared to unbiased experimental screens (all p < 0.0001, Fisher’s exact test). A negative log odds ratio indicates a depletion of positive interaction pairs in the literature. Notably, as we consider increasingly stringent p-value cutoffs to define significant genetic interactions, this log odds ratio becomes more negative, signifying that the underrepresentation of positive interaction pairs in the literature is more severe for highly significant genetic interactions. This underrepresentation has also persisted over all time periods examined (all p < 0.0001, Fisher’s exact test, Figure 2C), where each time period contains approximately one-third of the total literature in our sample (with unequal intervals reflecting uneven publication rates), indicating a consistent bias in research focus. These findings demonstrate that despite their scientific importance, positive genetic interactions are vastly underexplored in the scientific literature.

**Figure 2.**
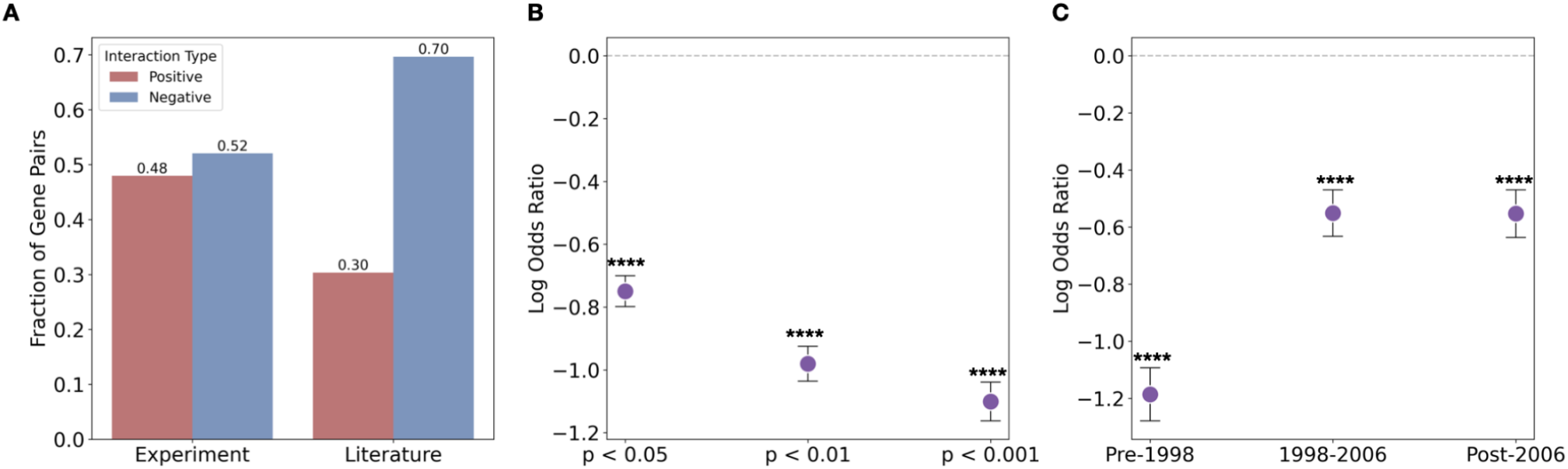
Positive interactions are under-represented in the literature. (A) Proportion of gene pairs with positive (red) versus negative (blue) interactions (using interactions with p < 0.05) among experimentally identified pairs (left) versus those co-studied in the literature (right). (B) Log odds ratio (literature vs. experiment) of positive to negative interactions at varying p-value thresholds; values below zero indicate depletion of positives. (C) The same log odds ratio over three publication periods, showing consistent under-representation of positive interactions. All error bars represent standard errors. Significance levels: NS, P ≥ 0.05; *P < 0.05; **P < 0.01; ***P < 0.001; ****P < 0.0001.

### The under-representation of positive interactions persists after unbiased interaction mapping

It is useful to distinguish two broad classes of explanation for this under-representation. The first is discoverability—physical and technical factors that make positive interactions harder to detect or measure, which an unbiased assay resolving interactions of both signs should alleviate. The second are social and practical factors that remain even once the existence of a genetic interaction is known, such as selection by investigators or reviewers over which gene pairs are studied, written up, and accepted for publication, and differential success by investigators in determining the causal basis of the observed interaction. While both pure discoverability and these various downstream investigatory factors may well be at play, we can gain insight into the role played by discoverability by taking into account the historical fact that the introduction of the SGA^7,8^ technology itself largely alleviated any issues due to differences in discoverability between positive and negative interactions. Thus, to probe the role of discoverability in the under-representation of positive genetic interactions, we conducted an event study asking how the reporting of positive versus negative interactions changed after the introduction of SGA, which could detect positive and negative interactions with equal facility, and which indeed provided researchers unbiased and comprehensive access to the sign and strength of interactions between all pairs of genes in the genome. In particular, to understand how the introduction of SGA changed researcher practice, we computed, in four-year bins, the log odds ratio of positive to negative interactions in the literature relative to their unbiased relative frequency among all genetic interactions (Figure 3).

**Figure 3.**
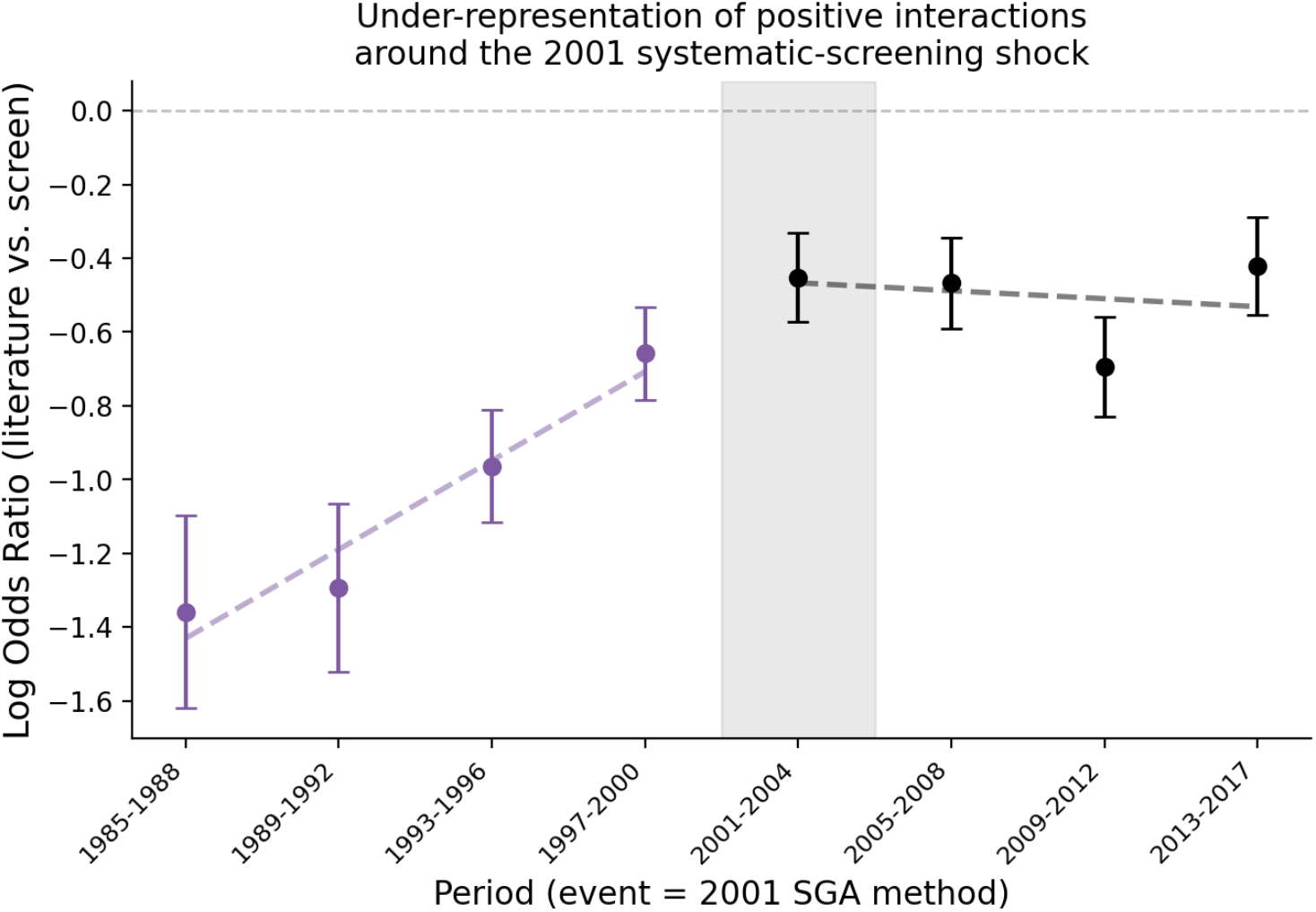
The under-representation of positive interactions persists after unbiased interaction mapping. In four-year bins (the final bin spans 2013–2017), the log odds ratio of positive to negative interactions in the literature relative to their unbiased relative frequency among all genetic interactions (higher values indicate less under-representation of positives). Points are bin estimates; dashed lines are the interrupted time-series fit before (purple) and after (black) 2001, and shading marks the bin in which the Synthetic Genetic Array was introduced. The deficit narrows steadily before ∼2001 (slope +0.24 per bin, p < 0.0001) but stops narrowing once positive and negative interactions become equally detectable (post-2001 slope −0.02, p = 0.65; change in slope −0.26, p < 0.0001). Error bars are one standard error.

Two features stand out. First, discoverability clearly mattered historically: before the early 2000s positive interactions were strongly under-reported, but this deficit was steadily shrinking (the log odds ratio rose from about −1.36 in 1985–1988 to −0.66 in 1997–2000), consistent with positive interactions becoming progressively easier to study. However, had limited discoverability been the binding constraint, a technology that detects positive interactions as readily as negative ones should, if anything, have accelerated this catch-up. Instead, the improvement halted: after the introduction of SGA in the early 2000s the log odds ratio plateaued near −0.5, with a post-2001 slope statistically indistinguishable from zero (−0.02, p = 0.65) and significantly flatter than the pre-2001 slope of +0.24 (change in slope −0.26, p < 0.0001; Figure 3). Because the deficit stopped closing around the point where detection could no longer plausibly be the binding constraint, its persistence is difficult to attribute to discoverability alone, and is consistent with social factors, such as selection by investigators or reviewers^19^ over which interactions are studied and reported, and/or practical differences in post-discovery investigatory success continuing to operate once any possible differences in discoverability were rendered irrelevant due to the introduction of comprehensive and unbiased measurements. We are cautious about this interpretation: the coincidence of the trend break with SGA does not by itself isolate the technology as the cause, and we cannot exclude a residual role for discoverability differences that resolved only gradually. What the data do indicate is that the persistence of the deficit past the point where detection ceased to be limiting is hard to explain by discoverability alone. We further note that the initial studies demonstrating the Synthetic Genetic Array illustrated the technique’s capabilities using synthetic lethality—that is, negative interactions—even though the same platform can assay positive interactions equally well^7^; this early choice of demonstration is itself consistent with the possibility that investigator preference contributes to the bias against studying positive interactions.

### Positive interactions seed more interconnected follow-on research

Finally, we asked whether the research that builds on positive-versus negative-interaction papers differs in quality or in structure. First, the downstream papers themselves are of slightly higher standing: ranking each citing paper within the citing literature of its publication year, the papers citing only positive interactions have slightly but significantly higher percentile citation rank and year-normalized citation impact than those citing only negative interactions (Figure 4A,B). Building on positive interactions therefore does not mean pursuing lower-impact directions—if anything, the opposite.

**Figure 4.**
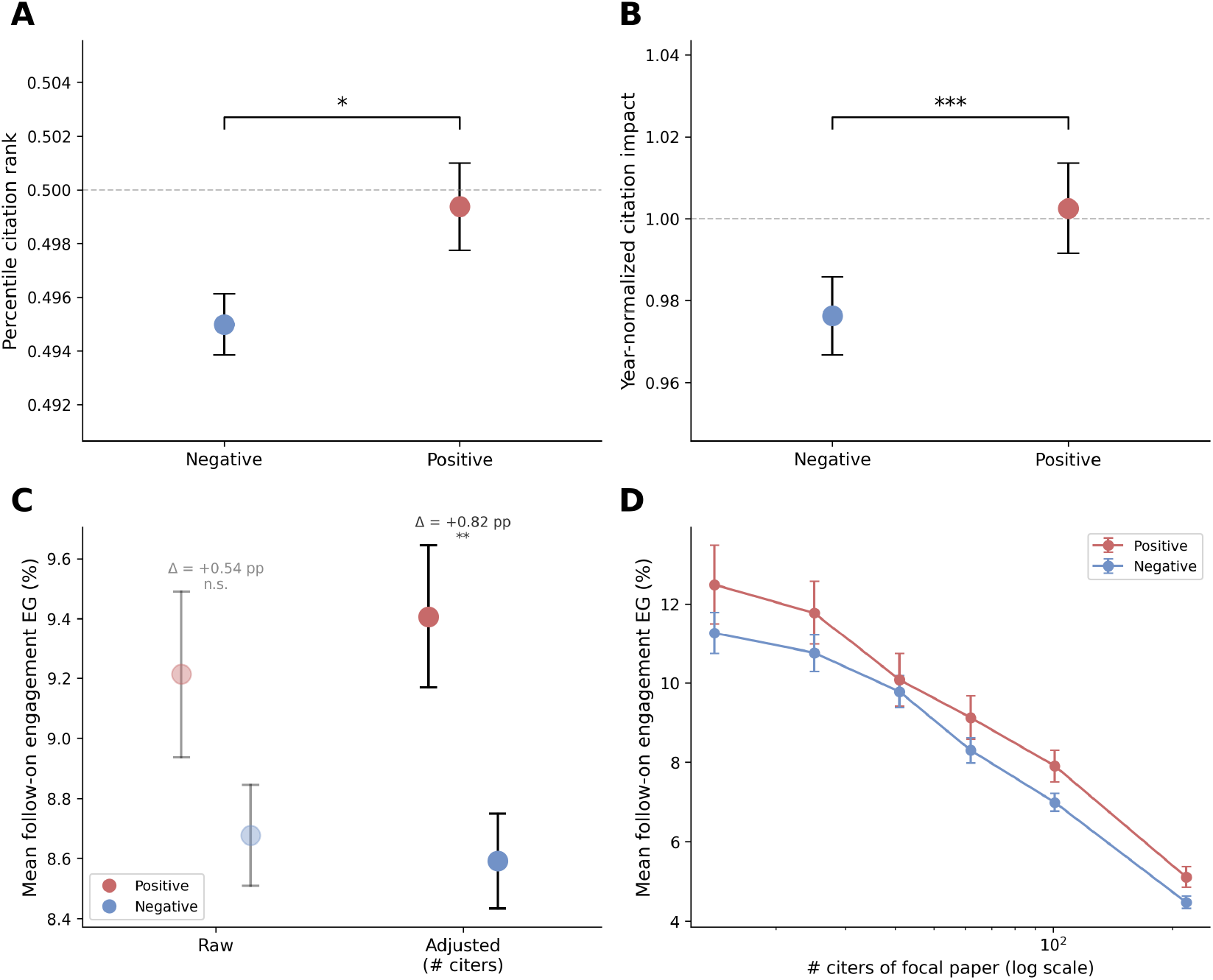
Downstream research building on positive-interaction papers is of slightly higher quality and more interconnected. (A, B) Citing papers ranked within the citing literature by publication year (disjoint citers that cite only one interaction class, one vote each): papers citing positive interactions have slightly but significantly higher (A) percentile citation rank and (B) year-normalized citation impact than those citing negative interactions. Axis labels, colors, dashed reference lines and standard-error bars follow Figure 1A,B. (C, D) Follow-on engagement (EG; the density of citation links among the papers that cite a focal paper, expressed as a percentage; see Methods; Hao et al.^20^): (C) raw group means (faded) overlap, but the citation-count-adjusted marginal means (solid) separate (+0.14 SD, p = 0.006); (D) mean EG within each citation-count stratum, positive ≥ negative in every stratum. Error bars are one standard error throughout. Significance (Mann–Whitney U in A, B; t-test on the sign coefficient in C): *P < 0.05, **P < 0.01, ***P < 0.001, ****P < 0.0001.

We then examined the structure of this literature. Following Hao et al.^20^, we measured follow-on engagement— the density of citation links among the papers that cite a given focal paper (as quantified by the EG metric, see Methods)—which captures whether the follow-on literature forms a connected line of subsequent research rather than a set of scattered, independent uses. Because larger citing sets tend to be less densely connected (the number of possible links grows quadratically with the number of citing papers) and positive-interaction papers are more cited, we compared positive and negative papers holding citation count fixed. Controlling for size in this manner, the follow-on literature of positive-interaction papers is significantly more interconnected (Figure 4C; +0.14 SD, p = 0.006). Examining the estimated mean engagement as an explicit function of citation count shows that positive-interaction papers have the higher estimated mean EG in every citation-count stratum (Figure 4D), the pooled size-adjusted estimate serving as the formal test. Thus positive interactions not only attract more follow-on work of slightly higher quality, that work more often coheres into a connected body of subsequent research, consistent with positive interactions opening new directions that the community then develops.

## Discussion

Previous studies have documented systematic patterns in how research attention is distributed across individual genes^21–26^. Here we extend this analysis to combinatorial innovation, long argued to be a dominant mode by which new ideas arise^10,27,28^, and consider the implications of which gene pairs are studied together. Our research reveals a significant, yet overlooked, phenomenon in genetic studies: positive genetic interactions, despite being linked to higher-impact publications, are consistently understudied. This observation directly challenges the conventional wisdom regarding which genetic interactions are most informative. It also resonates with established ideas in the sociology of science, which highlight the profound potential of unexpected discoveries.

Two further observations sharpen this picture. First, the under-representation of positive interactions does not simply reflect their being harder to detect: against a prior trend of steady improvement, the deficit stopped closing after the advent of unbiased high-throughput interaction mapping (Figure 3)—difficult to reconcile with a discoverability limitation alone—pointing to social factors, such as investigator and reviewer selection, or differences in the difficulty of explaining positive versus negative interaction, operating alongside any discoverability constraints. Second, the research built on positive interactions consists of papers with slightly higher per-paper citation standing and the resulting citation networks are more densely interconnected when conditioned on total citation count (Figure 4), suggesting that positive interactions more often open coherent new directions that the community develops, without a trade-off in downstream quality. The citation premium itself is robust to multivariate adjustment (Table S1) and to a non-GO novelty proxy.

We acknowledge a few limitations in our study. First, our reliance on literature curated by SGD might inadvertently favor highly cited papers. However, this potential bias should apply equally to both positive and negative genetic interactions, and thus not undermine our primary conclusions. Second, while citations are not a perfect metric of scientific impact due to social influences like prestige^13^—and, for readers less familiar with bibliometrics, we emphasize that our citation measures index the downstream attention and uptake a paper receives from the research community, not its intrinsic scientific value or any individual career outcome—there is no evident reason for such bias to disproportionately affect studies on positive versus negative interactions, making its impact on our results improbable. Finally, one could argue our findings are subject to survivorship bias, where only exceptionally strong positive interactions are studied, artificially inflating their perceived impact. Our data, however, indicate the opposite: negative interactions reported in the literature exhibit greater absolute strength (0.4452, 95% CI: [0.4295, 0.4609]) compared to positive ones (0.1196, 95% CI: [0.1134, 0.1257]); consistently, adjusting for interaction magnitude in the multivariate analysis (Table S1), if anything, strengthens the positive-interaction premium. This suggests that if positive interactions were studied at comparable strengths to negative interactions, their citation advantage would likely be even more pronounced.

More broadly, our study highlights the value of integrating sociological insights and “science-of-science”^29^ methodology to critically assess research strategies in biomedical science. Using bibliometric analyses, it becomes possible to empirically evaluate widely used research heuristics, such as the hypothesis that assays enhancing biological function yield more valuable insights than those that disrupt it^30^. We contend that developing this kind of meta-knowledge will be immensely beneficial to the biomedical community, fostering a more evidence-based approach to research design, and we suggest that cross-domain (positive) genetic interactions represent an under-exploited opportunity for discovery.

## Acknowledgements

This work was supported by NIH grant R35 GM133613 (MS, DMM), as well as additional funding from the Simons Center for Quantitative Biology at CSHL (MS, DMM).

## Data availability

All code and data necessary to reproduce the analyses in the manuscript will be published as a github repository upon publication.

## Materials and Methods

### Data

Data for this study were drawn from several sources. Genetic interaction data were obtained from the Costanzo et al. (2016) large-scale screen (https://boonelab.ccbr.utoronto.ca/supplement/costanzo2016/), focusing on 1,331,306 significant non-essential gene pairs (p < 0.05). This selection prioritized interaction scores derived from knockout experiments, considered the gold standard, and avoided trivial positive interactions often associated with essential genes^31^. Gene literature data were acquired from the Saccharomyces Genome Database (SGD) gene literature database (http://sgd-archive.yeastgenome.org/curation/literature/,gene_literature.tab file). This systematically curated database links 418,111 papers to their associated yeast genes. For our analysis, we first counted the number of genes associated with each paper, then retained only those papers linked to precisely two genes, allowing us to attribute scientific impact to specific gene pairs. Bibliometric data were sourced from the iCite database (https://nih.figshare.com/authors/iCite/7026758) and matched to the SGD gene-pair literature using PubMed IDs (PMIDs), filtering for ‘research articles’ as defined by iCite, resulting in 1,981 papers with citation data associated with gene pairs involved in significant genetic interactions. For the additional analyses introduced in this revision, we also used: paper subfields and topics from OpenAlex^17^; biological pathway memberships from the Kyoto Encyclopedia of Genes and Genomes (KEGG)^18^; and the paper-to-paper citation network from the NIH Open Citation Collection^12^ (used to compute follow-on engagement).

### Scientific Impact Metrics

Scientific impact was assessed using two different year-normalized metrics. The primary metric was year-specific percentile citation rank, defined as the percentile of a paper’s citation count within all gene literature articles published in the same year that could be linked to iCite. This method provides fair comparisons across publication years, accounting for citation accumulation over time. The alternative metric, year-normalized citation impact, was calculated by dividing a paper’s citation count by the average citation count for papers published in the same year.

### Novelty Mediation Analysis

To assess whether the higher impact of positive genetic interactions is driven by their novelty—specifically, their tendency to connect functionally distant genes—we computed GO-term similarity for each gene pair using the GOATOOLS library^32^. Annotations from the Saccharomyces Genome Database GAF file were propagated to all parent terms, and for each pair we averaged all pairwise similarities under two complementary metrics: Wang’s graph-based measure of structural overlap in the GO hierarchy, and Lin’s information-content measure emphasizing term specificity. To isolate novelty’s effect, we paired every positive interaction with a negative interaction by calculating the Mahalanobis distance between their (Wang, Lin) similarity vectors and solving the resulting assignment problem to minimize overall distance, yielding one-to-one matched sets without discarding any positives. We also created a size-matched random control by sampling negatives equal in number to the positives. As a non-GO check, we repeated the matching using KEGG pathway co-membership (the Jaccard overlap of the two genes’ pathways) as the measure of functional similarity, which likewise rendered the citation-rank advantage non-significant after matching.

### Multivariate Regression

As a complement to matching, we regressed (Table S1) each impact metric on an indicator function for positive genetic interaction sign together with covariates, fitting a four-model ladder per outcome: unadjusted; adding the absolute interaction magnitude |I|; adding number of authors, journal venue, publication-year (decade) indicators, and the paper’s OpenAlex primary-topic subfield; and a fully saturated model combining all of these covariates with |I|. Models were fit by ordinary least squares with HC3 heteroskedasticity-robust standard errors. We use the OpenAlex subfield—derived from a paper’s text and citations and therefore independent of the gene-level GO annotations—as a field-composition control; controlling instead for GO-derived subfield categories absorbs the premium, as expected, because those categories proxy the novelty mediator itself.

### Temporal Analysis of Reporting Bias

To test whether the under-representation of positive interactions changed once an unbiased assay became available (Figure 3), we binned the literature into four-year periods (the final bin, 2013–2017, spans five years) and, for each bin, computed the log odds ratio of positive to negative interactions relative to their unbiased relative frequency in the Costanzo screen^2^. We summarized the temporal trend with an interrupted time-series fit allowing the slope to change at the introduction of the Synthetic Genetic Array^7^, taking 2001 as the event.

### Follow-on Engagement

To characterize the structure of downstream research (Figure 4), we adopted the follow-on engagement metric of Hao et al.^20^, which measures how densely a paper’s citing papers cite one another. For a focal paper with n citing papers, engagement is EG = 2k / [n(n − 1)], where k is the number of citation links among those n citing papers. Because citations run from later to earlier work, this subgraph is acyclic and each connected pair of citers contributes exactly one link, so k is the number of connected pairs and EG = k / [n(n − 1)/2] is the fraction of the n(n − 1)/2 possible pairs of citers that are directly linked, expressed as a percentage. The citer-to-citer links were obtained from the NIH Open Citation Collection^12^. Because EG tends to decrease with n (the number of possible pairs grows quadratically), we compared positive and negative interaction papers both raw and controlling for citation count (log number of citers) by ordinary least squares with HC3 robust standard errors, and verified the difference within citation-count strata. We restricted this analysis to focal papers with at least 10 citing papers (1,714 of the 1,981 focal papers; 526 positive and 1,188 negative), because EG is a density over the n(n − 1)/2 possible pairs of citers and is too coarse to be informative below that: among focal papers with between two and nine citers, a third take a value of exactly 0 or exactly 100. The comparison is not sensitive to this cutoff— lowering the threshold to five citers or raising it to twenty leaves the citation-count-adjusted advantage of positive-interaction papers positive and statistically significant (+0.10 to +0.14 SD, p < 0.05).

## Supplementary Materials

Supplementary Table S1 reports the multivariate robustness of the positive-interaction citation premium across a control ladder, supporting the impact results in the main text. (The non-GO pathway-novelty mediation check is summarized in the main text.)

**Table S1.**
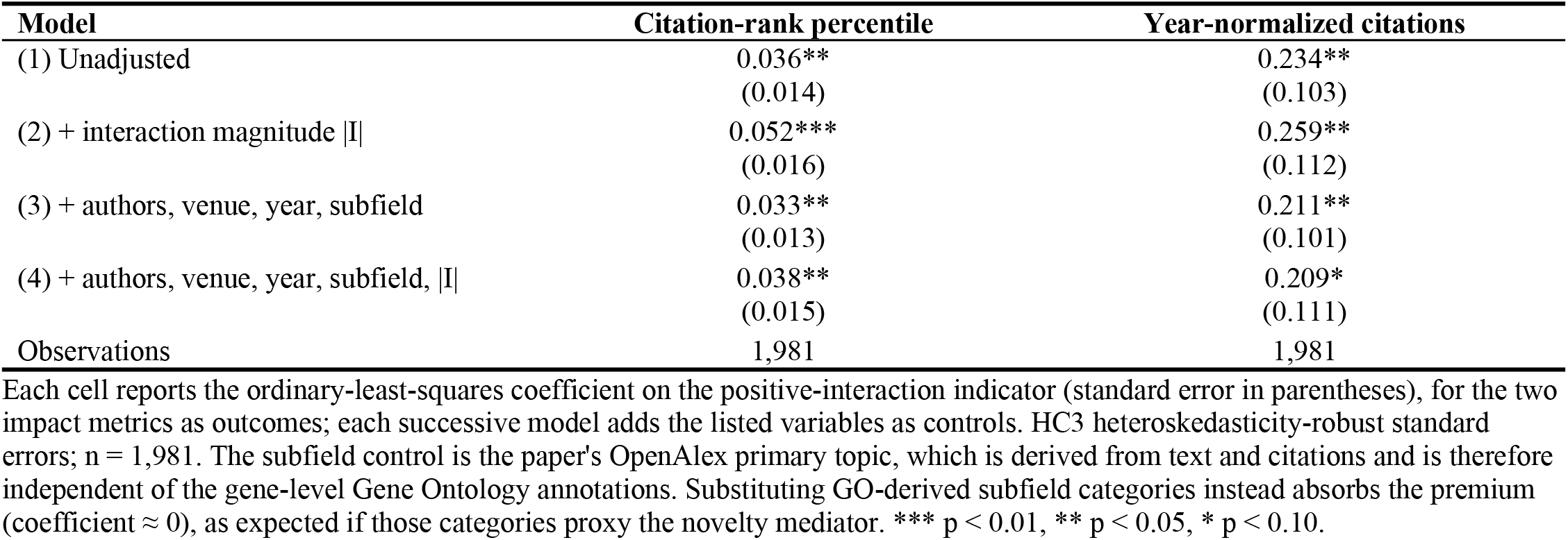
Multivariate robustness of the positive-interaction citation premium.

